# Self-Supervised Behavioral Representations Across the Life Course: A Killifish Case Study

**DOI:** 10.64898/2026.06.23.733896

**Authors:** Jen-Chien Chang, Teruhisa S. Komatsu, Shuichi Onami

## Abstract

Self-supervised foundation models of aging are increasingly built from longitudinal data (biobanks, electronic health records, wearables) that is inherently incomplete: no individual is followed across a whole lifetime, and how much of each life is captured varies widely. This raises two linked questions: is it worth modeling an individual’s whole life course rather than its current state, and can such a model be built from brief, fragmentary records? No human cohort can settle them, because none offers a complete life to compare against. We turn to the African turquoise killifish (*Nothobranchius furzeri*), tracked from youth to natural death in publicly released recordings, as a controlled testbed: its complete lifespans provide the full-life reference that human data lacks. On these data we build LifeMAE, a two-stage self-supervised model: a *day encoder* that summarizes each day of behavior, then a *life-course encoder* over the trajectory of those daily summaries. We find that the day encoder alone is already strong: from a single day of behavior it predicts chronological age, separates long-from short-lived individuals (coarsely), and flags nearness to death. Adding the life-course encoder improves on none of the three; each is matched by trivially aggregating the day-level predictions (a smoother for age, an early-life average for lifespan). Near-term mortality seems the exception, where the whole-life model looks far better (AUROC 0.81 to 0.91), but the gain is not behavioral: it reflects where each day falls within the observation window (a cue supplied by the model’s encoding of time), and a single-day model given that cue closes the gap at any observation length. For these traits, an individual’s place in its life course is legible from a single day: the trajectory stage is unnecessary, and the record it needs is as short as one day, the finest grain our day-level setup resolves. For characterizing a cohort, this favors observing many individuals briefly over tracking a few for long. The result joins a growing body of work in which deep and foundation models, fairly benchmarked, fail to beat deliberately simple baselines. We add a concrete mechanism for the over-optimism: a model’s encoding of time can leak the very quantity it predicts, which backward-looking evaluation mistakes for learned biology, so only evaluation fixed to the moment of prediction is trustworthy.

## 1 Introduction

Aging is a fundamentally longitudinal process: the same individual must be followed over time for the trajectory of decline to become visible. Dense, continuous behavioral monitoring is an attractive substrate for studying it, because spontaneous behavior is cheap to record and integrates neural, muscular, metabolic, and motivational state into a single non-invasive readout [Datta et al., 2019]. Coupled with self-supervised representation learning, such recordings promise label-light biomarkers of biological aging built from behavior alone.

### The problem: real longitudinal data is fragmentary

The data on which such models would ultimately be trained (human biobanks, electronic health records, and wearable-sensor cohorts) share a structural limitation: they are *partial-life*. No participant is observed continuously from adolescence to death; each contributes a short, arbitrarily placed slice of a long life (a week of wrist accelerometry, sporadic clinical visits), and the fraction of the lifespan captured is neither controlled nor known. Recent longitudinal foundation models built on such data (Transformer models of electronic health records [Li et al., 2020] and large-scale generative models of lifelong health trajectories such as Delphi-2M [Shmatko et al., 2025] and LifeClock [Wang et al., 2025a]) show that useful representations can be learned from fragmentary trajectories. But they cannot say which capabilities actually require the longitudinal trajectory, nor how much they suffer under fragmentation, because there is no full-life ground truth to ablate against: one never observes the complete trajectory that the fragments are drawn from.

### A controlled testbed

We turn to a model organism where full-life ground truth does exist. The African turquoise killifish (*Nothobranchius furzeri*) is the shortest-lived vertebrate bred in captivity, with a median lifespan of four to six months and well-characterized aging phenotypes [Harel et al., 2015]. Bedbrook et al. [2026] recently released continuous behavioral tracking of hundreds of killifish from adolescence to natural death, processed into a per-day vocabulary of behavioral “syllables”: a rare public dataset combining hundreds of individuals, daily behavioral resolution, and complete lifespan coverage in a vertebrate. We can therefore train on the complete trajectories, treat that as an upper bound, and systematically degrade the training data into episodic and partial-life regimes that mimic biobank-style sampling, measuring exactly what is lost.

### Approach

We build **LifeMAE**, a two-stage self-supervised *life-course masked autoencoder*. A *day encoder* (a BERT-style masked autoencoder [Devlin et al., 2019, He et al., 2022] over 10-minute behavioral-state histograms) learns a representation of a single day of behavior. A *life-course encoder* (a Transformer over the age-ordered sequence of frozen day embeddings) learns a representation of the individual’s trajectory. Holding the day encoder fixed, we retrain the life-course encoder under regimes that vary how much of each individual’s life is seen (from complete lifespans down to disjoint 3-day episodes), read it out across observation windows from a few days to the whole life, and ask a sharp question of every target: does the life-course stage add anything over the day-level representation it is built on? This subsumes two questions in one: whether an individual’s *trajectory* can be learned from fragments, and how little observation is needed to *place* an individual in its life course. We find the first premise empty for every trait we can ground-truth, which collapses the question to the second; its answer is a single day.

### Contributions

#### 1. A single self-supervised day encoder recovers the published aging read-outs

From one day of behavior, light probes read chronological age (*R*^2^ = 0.88) and proximity to death (AUROC 0.81); from a short early-life window of those single-day representations, eventual lifespan (AUROC 0.98). These match or beat a train-only refit of the dataset’s own factor model: a single, label-free substrate behind every read-out (Section 4.1).

#### 2. Holding that day encoder fixed, the life-course stage adds nothing

Because the day encoder is frozen and only the life-course encoder is retrained, any difference is attributable to the trajectory stage alone; under that control, every trait we can ground-truth is matched by the single day plus a *trivial, non-learned* operation: a temporal smoother for age, an early-life pool for lifespan, and a within-window timing cue for 30-day mortality. Mortality is the subtle case: the life-course stage appears to add a large gain (AUROC 0.81 → 0.91), but it is supplied by the model’s encoding of time, not by behavior, and vanishes once a single-day probe is given the same timing information (the residual is statistically indistinguishable from zero at every observation window; Sections 4.2 and 4.3). We read this as *cross-sectional sufficiency*: an individual’s place in its life course is legible from a snapshot.

#### 3. An evaluation trap explains why the life course looked essential

Positional encodings leak task-specific position: absolute age inflates age estimates (the model reads the target off its own input), and window-relative age inflates retrospective mortality (within-window position tracks remaining life). Ordinary centered-window probing mistakes this for learned dynamics; only evaluation fixed to the moment of prediction is confound-free. With probe expressiveness and fish-level multi-seed intervals, these choices change the conclusions and transfer to other life-course models (Section 4.4).

### Scope

This is deliberately a single-species study: the killifish is the instrument, chosen because it uniquely supplies the full-life upper bound the question requires, not as an end in itself; we make no cross-species claims. Our readouts are three traits with hard full-life ground truth, and our claim is about *characterizing* an individual’s current state, not forecasting its future: the deployable, forward-looking mortality effect is weak, and we say so (Section 6). Traits that genuinely turn on how behavior changes over time (a rate of aging, a change point) remain an open and reachable target; we simply find none among the traits we can presently ground-truth. We present this as a controlled characterization, against a complete-life reference, of when self-supervised life-course representations help and whether they can be built from fragmentary behavioral data, a question the human longitudinal foundation models that motivate this work cannot settle from their own data. We discuss the implications for biobank-scale design in Section 5.

## 2 Related Work

### Longitudinal and biobank foundation models

A growing body of work learns representations of individual health trajectories from large longitudinal datasets: Transformer models of electronic health records [Li et al., 2020, Rasmy et al., 2021, Choi et al., 2020] and, more recently, generative models of lifelong disease and aging trajectories at biobank scale, such as Delphi-2M [Shmatko et al., 2025] and LifeClock [Wang et al., 2025a]; and, closest to our own setting, self-supervised models of wrist-worn accelerometry that learn human activity and health representations from short, free-living wear windows [Yuan et al., 2024, Xu et al., 2025], the human analog of the behavioral records we study. These models establish, operationally, that useful representations can be learned from fragmentary records: no individual is observed across a full lifespan, and each contributes only a sparse, arbitrarily placed slice. What they cannot provide is a controlled account of which capabilities actually require the longitudinal trajectory, and how they degrade under fragmentation, precisely because the full trajectories the fragments are drawn from are never observed. Our study supplies that missing control, on a model organism whose complete trajectories are observed. This caution echoes a broader reckoning in single-cell genomics, where critical benchmarking has repeatedly found foundation models no better than deliberately simple baselines for perturbation-effect prediction [Ahlmann-Eltze et al., 2025].

### Self-supervised behavioral phenotyping

Unsupervised discovery of behavioral “syllables” from video (pioneered by MoSeq [Wiltschko et al., 2015] and built on pose trackers such as DeepLabCut [Mathis et al., 2018] and SLEAP [Pereira et al., 2022], with state-space successors like Keypoint-MoSeq [Weinreb et al., 2024]) turns raw video into discrete behavioral-state time series. Bedbrook et al. [2026] apply a Gaussian hidden Markov model to killifish pose dynamics to obtain a 100-state syllable vocabulary, then summarize each day with non-negative tensor component analysis [Williams et al., 2018]. We consume the same per-day syllable histograms but replace the fixed factor-analytic day-level summary with a learned encoder, so that downstream differences are attributable to the representation rather than to the upstream tokenizer.

### Masked self-supervised modeling

Our encoders instantiate the masked-prediction recipe that has proven effective across modalities: masked language modeling [Devlin et al., 2019], masked image modeling [He et al., 2022, Xie et al., 2022], and masked modeling of time series [Zerveas et al., 2021, Nie et al., 2023]. The day encoder is a masked autoencoder over a day of behavioral-state histograms; the life-course encoder applies the same principle one timescale up, masking whole days within an individual’s trajectory. Two design choices adapt the recipe to behavior: cyclic positional encoding that bakes circadian periodicity into the geometry, and continuous-span masking, since adjacent 10-minute bins are strongly autocorrelated and single-bin masking is trivially solved by interpolation [Joshi et al., 2020]. Beyond masked modeling, contrastive self-supervised time-series methods such as TS2Vec [Yue et al., 2022] and TS-TCC [Eldele et al., 2021] offer alternative trajectory encoders; because our question concerns what a life-course representation captures rather than which temporal architecture is best, we leave a head-to-head comparison of encoders to future work (Section 6).

### Lifespan-scale behavioral representation

Modeling behavior at the scale of whole lifetimes, rather than single recording sessions, is comparatively unexplored. The lifelong *C. elegans* work of Stroustrup et al. [2016] tracked behavior to death but relied on bulk activity statistics rather than learned trajectory representations. A growing family of self-supervised behavioral models learns rich motion representations from pose or video (variational motion embeddings [Luxem et al., 2022], multi-task and contrastive objectives [Azabou et al., 2023, Schneider et al., 2023], and, most directly related to ours, (hierarchical) masked autoencoders over pose trajectories and video [Stoffl et al., 2024, Wang et al., 2025b]) but they represent behavior *within* single continuous recordings, typically a minute to tens of minutes long (across multiple within-recording timescales, for the hierarchical and multi-task variants); none are organized around an individual’s behavior across its *lifespan*, with age as the temporal axis, nor do they address building such a representation from fragmentary, multi-day data. Our life-course encoder is explicitly an individual-trajectory model: its representation is conditioned on an individual’s behavioral trajectory to date, which is what makes the partial-life question (whether, and for which targets, that trajectory is actually needed) well posed.

## 3 Methods

Figure 1 summarizes the two-stage architecture of LifeMAE.

**Figure 1.**
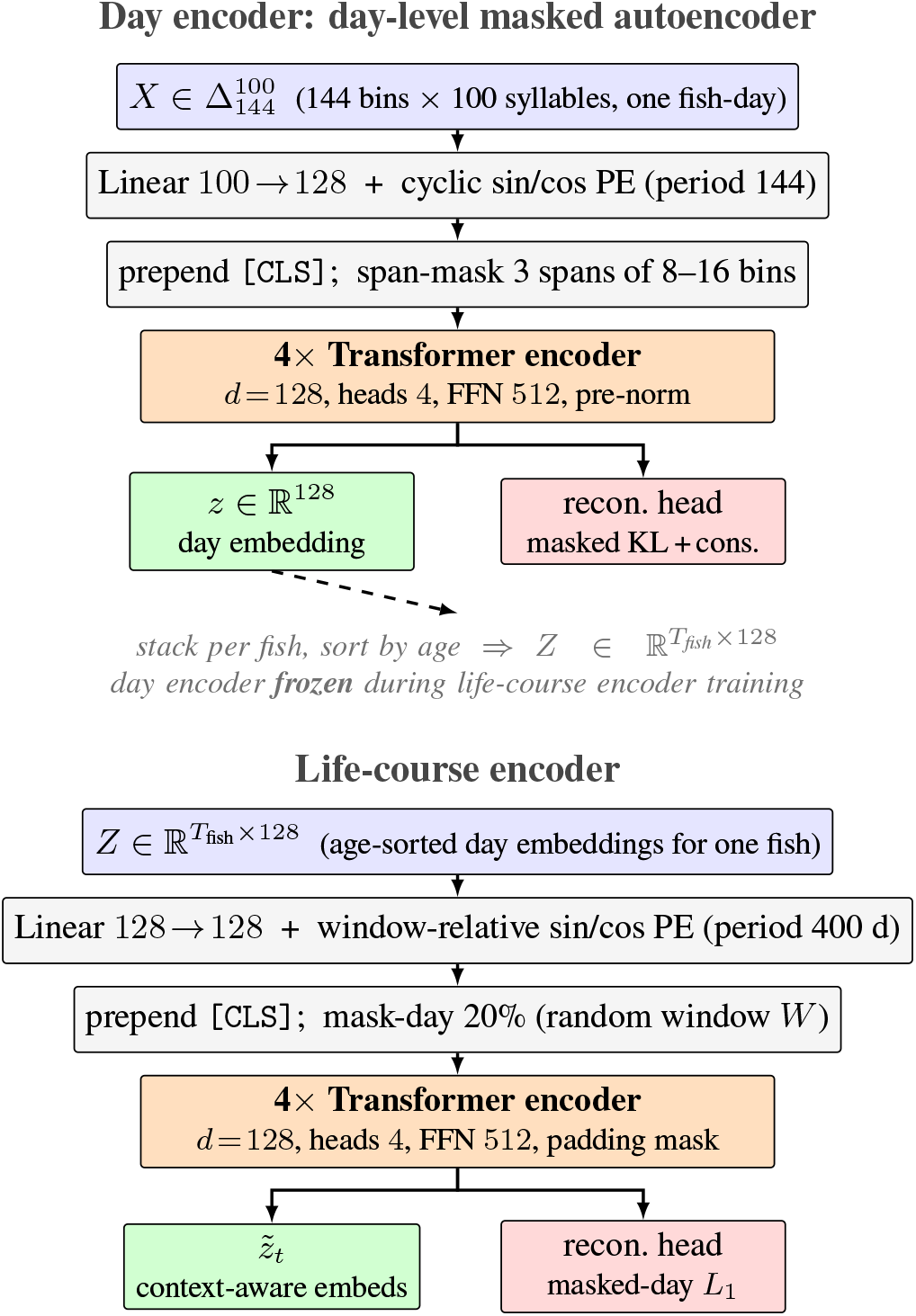
Two-stage architecture of LifeMAE (day encoder and life-course encoder). **Day encoder** (top) is a BERT-style masked autoencoder over 10-minute syllable histograms for one fish-day at a time: a cyclic sin/cos positional encoding with period 144 bakes circadian periodicity into the geometry; span masking (3 spans of 8–16 bins) forces the model to use broad circadian context; the `[CLS]` output is the 128-d day embedding and the pretext task is bin-level reconstruction (masked KL plus a small consistency term). **Life-course encoder** (bottom) is a second Transformer over the age-sorted sequence of *frozen* day embeddings for each fish, with *window-relative* sin/cos positional encoding (period 400 d) on each day’s age relative to the window start, so that recording gaps advance the encoding by the correct number of days while absolute age is withheld. It is trained purely by masked-day smooth-*L*_1_ reconstruction (20% of days held out per window), yielding context-aware per-day embeddings 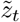 that are frozen for downstream probes. Each encoder has ∼ 0.8M parameters; only the life-course encoder receives gradients during its training, while the day encoder stays frozen.

### 3.1 Data and preprocessing

We use the public Zenodo release of Bedbrook et al. [2026] (CC-BY-4.0; DOI in *Data and code availability*): continuous video-derived behavioral-state assignments for 234 African turquoise killifish tracked individually from puberty to death. After light quality filtering we retain *N*_fish-days_ = 24,277 recording days across the 234 fish, of which 116 end in natural death (the remainder are censored at sacrifice or removal). Each *fish-day* is a matrix 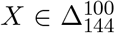 ten-minute bins, each a probability distribution over 100 hidden-Markov-model behavioral “syllables.”

#### Splits and leakage control

All splits are disjoint at the level of individual fish (`full_fish_name`); days from one fish never span two splits. Our primary split is a fish-held-out 70/15/15 partition (164 train fish), assigned at random over fish. Every preprocessing statistic that is fit-and-applied (feature scalers, missing-bin fills, and the tensor-factorization baseline) is fit on training fish only, and survival-derived fields (`lifespan`, `status`, remaining life) are never used as encoder inputs. Table 1 reports cohort sizes by split.

**Table 1.** Cohort sizes by split (fish-held-out 70/15/15). The 30-day-mortality probe is restricted to naturally-dying fish, where the label (death within 30 days) is well defined; “30-d+ days” counts fish-days with a positive label, with the per-day positive rate in parentheses. The held-out mortality cohort is small (21 natural-death test fish), which is why we report fish-level bootstrap intervals throughout.

| Split | Fish | Fish-days | Natural-death fish | Natural-death days | 30-d+ days (rate) |
| --- | --- | --- | --- | --- | --- |
| Train | 164 | 16,015 | 81 | 10,462 | 2,217 (0.21) |
| Val | 35 | 4,311 | 14 | 2,684 | 392 (0.15) |
| Test | 35 | 3,951 | 21 | 3,245 | 592 (0.18) |
| Total | 234 | 24,277 | 116 | 16,391 | 3,201 (0.20) |

### 3.2 Day encoder

The day encoder is a BERT-style [Devlin et al., 2019] bidirectional Transformer over one *fish-day*. Each fish-day is projected token-wise to the model dimension, positional-encoded, and passed through 4 encoder layers (*d* = 128, 4 heads, feed-forward 512, pre-norm, dropout 0.1); a learned `[CLS]` token yields the 128-dimensional day embedding, and a shared linear head reconstructs the 100-d syllable distribution at each of the 144 positions. The encoder has ∼0.8 M parameters.

#### Cyclic circadian position encoding

We use fixed sin/cos features with period equal to the full day,

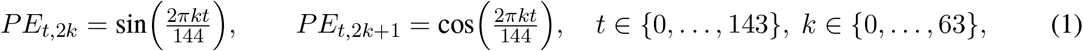

so that bin 143 (23:50) is adjacent to bin 0 (00:00): circadian periodicity is encoded in the geometry rather than learned from data.

#### Continuous-span masking and objective

Because syllable composition is strongly autocorrelated across adjacent bins, we mask 3 contiguous spans of 8–16 bins each (≈23% of the day on average, up to 33%) rather than isolated bins, and predict the original distribution only at masked positions. With 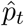 the predicted and *p*_*t*_ the true distribution at bin *t* and *M* the masked set,

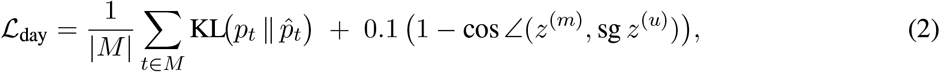

where the second term is a mild mask-invariance regularizer: the cosine distance between the `[CLS]` embedding of a masked view and the stop-gradient [Chen and He, 2021] `[CLS]` embedding of the unmasked day. The direction 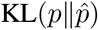 avoids penalizing the model for placing mass on syllables the target assigns zero probability.

#### Training

Short-lived fish are over-represented at young ages, so an age-balanced sampler draws uniformly over (10-day age-bucket × fish) pairs. We train with AdamW (lr 3 × 10^−4^, weight decay 0.01), a cosine schedule, and mixed precision; the day encoder is then frozen.

### 3.3 Life-course encoder

The life-course encoder is a second Transformer over a fish’s age-ordered sequence of frozen day embeddings *z*_1_, …, *z*_*T*_ ; it never re-reads the raw syllable tensor. It uses the same 4-layer, *d* = 128 backbone, with sinusoidal positional encoding of *elapsed time within the observed window*: each day’s age is taken relative to the first day of the window, Δ*a*_*t*_ = *a*_*t*_ − *a*_1_,

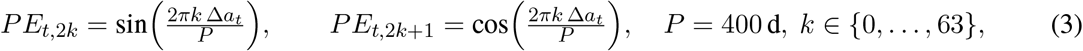

so a 14-day recording gap still advances the encoding by 14 days (irregular sampling is respected) while the *absolute* age is withheld. This choice is deliberate: because chronological age is itself a downstream read-out target, encoding absolute age would inject that target into the representation and inflate the apparent value of the life-course stage; a window-relative encoding keeps trajectory order and inter-day gaps while concealing how old the fish is. We quantify the difference against an absolute-age encoding in Section 4.4. The encoder is trained purely by masked-day reconstruction: 20% of the days in each sampled window are replaced with a learned mask token and reconstructed with a smooth-*L*_1_ loss on the 128-d embedding. To create trajectory diversity from 164 training fish, each training step samples a random contiguous window per fish (length drawn uniformly from [min(14, *T*), *T*], padded to a fixed maximum); the lower bound min(14, *T*) lets a fragmentation episode shorter than 14 days, such as `fixed-3`, contribute its whole length. The encoder emits context-aware per-day embeddings 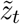 and a fish-level `[CLS]` pool, has ∼ 0.8 M parameters, and is trained with AdamW (lr 10^−4^). Each read-out uses a local window, but the encoder is trained on windows drawn from across the lifespan and pooled over the cohort, so it learns to represent any life stage; this pooled-trajectory training, not any single long input, is what makes it a life-course encoder.

### 3.4 Episodic and partial-life training regimes

The fragmentation experiment fragments the life-course stage while holding the day encoder fixed: the day-level foundation is trained once on all training fish-days and reused unchanged across regimes, so it cancels in the comparison and the ablation isolates trajectory aggregation. For each regime, an `episode_spec` deterministically truncates each training fish’s available days before window sampling:

`full`

every training fish contributes its complete trajectory (the upper bound).

`fixed-`*L*

each fish contributes a single *L*-day contiguous episode (*L* ∈ {3, 7, 14, 28, 56}), placed uniformly at random; `fixed-3` is the most severe fragmentation.

Each trained encoder is then read out across a range of inference-context lengths *L*_inf_ without further retraining, so that training- and inference-time coverage are varied independently over the (training-window × inference-window) surface.

### 3.5 Evaluation

Embeddings from both encoders are frozen and read out by simple probes fit on training fish, selected on validation fish, and reported on test fish; all features are standardized (statistics fit on training fish). To separate representation quality from probe capacity we report two probe levels: **Level A**, a linear probe (ridge for age, logistic for classification); and **Level B**, a small two-layer MLP head (Linear → GELU → Linear). We evaluate age regression (mean and median absolute error, *R*^2^), long-versus-short lifespan classification (AUROC, threshold at the train-fish median lifespan, naturally-dying fish only, where the lifespan is known), behavioral life-stage agreement (*k*-means with *k*=6 against the released six-stage labels; adjusted Rand index), and a 30-day mortality probe (AUROC), restricted to the 116 naturally-dying fish so that the label (death within the next 30 days) is well defined.

#### Inference protocol and baseline

At inference an encoder is given an *L*_inf_-day window of a test fish and read out per day. Our primary protocol is *retrospective*: the window is a contiguous slice of the observed trajectory and each day is encoded with bidirectional context, so we characterize the recorded life against the full-life upper bound rather than forecast forward. As a deployability check we also report a *causal* protocol, a trailing window whose final day is the target, so no future days are visible. The comparison baseline uses the frozen day embeddings without the life-course encoder, read per day in the retrospective protocol (so the gap isolates the value of trajectory context) and mean-pooled over the trailing window in the causal protocol, through the identical probe, so differences reflect trajectory modeling rather than the day representation.

#### Multi-seed paired evaluation

Encoder-initialization variance is non-negligible relative to the effects we measure, so every reported comparison aggregates a 3 × 3 factorial of day-encoder and life-course-encoder seeds (nine cells), and confidence intervals are a paired bootstrap that resamples test *fish* (not fish-days), nested over the nine seed cells. We report the paired gap between the life-course representation and the day-level baseline for both chronological age (median absolute error) and 30-day mortality (AUROC), at matched inference context.

#### Within-window position control

The window-relative encoding makes each day’s position within the observed window available to the model, and that position can itself track the outcome: for a short-lived fish a centered month covers most of its life, so a day late in the window is close to death. We therefore test whether the life-course mortality gain is positional rather than behavioral. We form the same sinusoidal within-window position vector the encoder receives, PE(Δ*a*_*t*_) of Eq. (3), and read mortality through the identical probe from three feature sets: the day embedding alone (behavior), the position vector alone (no behavior), and their concatenation (behavior + position). If the life-course stage adds genuine trajectory structure it should beat behavior + position; if its gain is the position cue, the two should match. We additionally concatenate a window-mean of the day embeddings (behavior + position + pooling), and, for the per-fish lifespan target, where a fish-level read-out is natural, compare the life-course `[CLS]` embedding against a plain mean-pool of the day embeddings, to check that a learned fish-level summary does not rescue the trajectory stage where the per-day read-out does not.

#### Learned-representation comparator (TCA)

As a learned-representation comparator we refit the non-negative tensor component analysis of Williams et al. [2018], Bedbrook et al. [2026] (rank 45) on the training fish only, and read it out through the identical standardized probes, so that differences reflect the representation rather than the probe or the preprocessing.

## 4 Results

We first establish that the learned day representation captures real aging signal (Section 4.1). We then ask, for every target, whether the life-course stage adds anything over that day-level representation, and find that it does not: chronological age and eventual lifespan are already legible day by day, and the one apparent exception, 30-day mortality, dissolves under control (Section 4.2). We trace that apparent exception to a within-window position cue introduced by how the encoder represents time, not to learned trajectory structure (Section 4.3). We close with the evaluation choices that produced these measurements, and that we expect transfer to other longitudinal-representation studies (Section 4.4).

### 4.1 The day representation is biologically meaningful

Under strict fish-held-out evaluation, a single day of behavior encoded by the day encoder predicts chronological age with a median absolute error of 15.7 days (mean 21.6; *R*^2^ = 0.84) under a linear probe, improving to 12.6 days (*R*^2^ = 0.88) under a small nonlinear probe (Table 2). The same representation, mean-pooled over a short early-life window, separates long-from short-lived individuals (naturally-dying fish, train-fish median split; AUROC 0.98, 95% fish-level CI [0.91, 1.00]; partly age-correlated and threshold-sensitive at this sample size, Section 6), agrees with the released six-stage behavioral life-stage labels (*k*-means adjusted Rand index 0.45), and supports a 30-day mortality probe (AUROC 0.81 [0.74, 0.88]). These match or exceed a train-only refit of the dataset’s own non-negative tensor-factorization representation: refit on training fish and read through the identical standardized probes, that baseline reaches age *R*^2^ of 0.75 (linear) and 0.73 (nonlinear), with median errors near 20 days, below the learned day encoder at both probe levels. Projected to two dimensions, the day embeddings organize along biologically meaningful axes: age, lifespan, and feeding regime (Fig. 2), so the representation is a faithful behavioral substrate, not a compression artifact. Notably, the day encoder never sees an individual’s trajectory: it is trained on single days pooled across the cohort, so this age, lifespan, and life-stage structure is learned from cross-sectional behavior alone. This single-day representation is the strong baseline that the rest of the paper asks the life-course stage to beat.

**Table 2.**
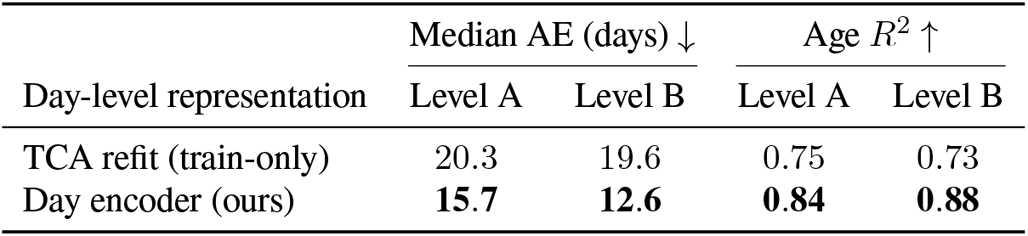
The day-encoder representation is biologically meaningful. Per-fish-day chronological-age prediction (strict fish-held-out, standardized features; median absolute error and coefficient of determination) for the train-only tensor-factorization refit and the learned day encoder, at a linear (Level A) and a small nonlinear (Level B) probe.

**Figure 2.**
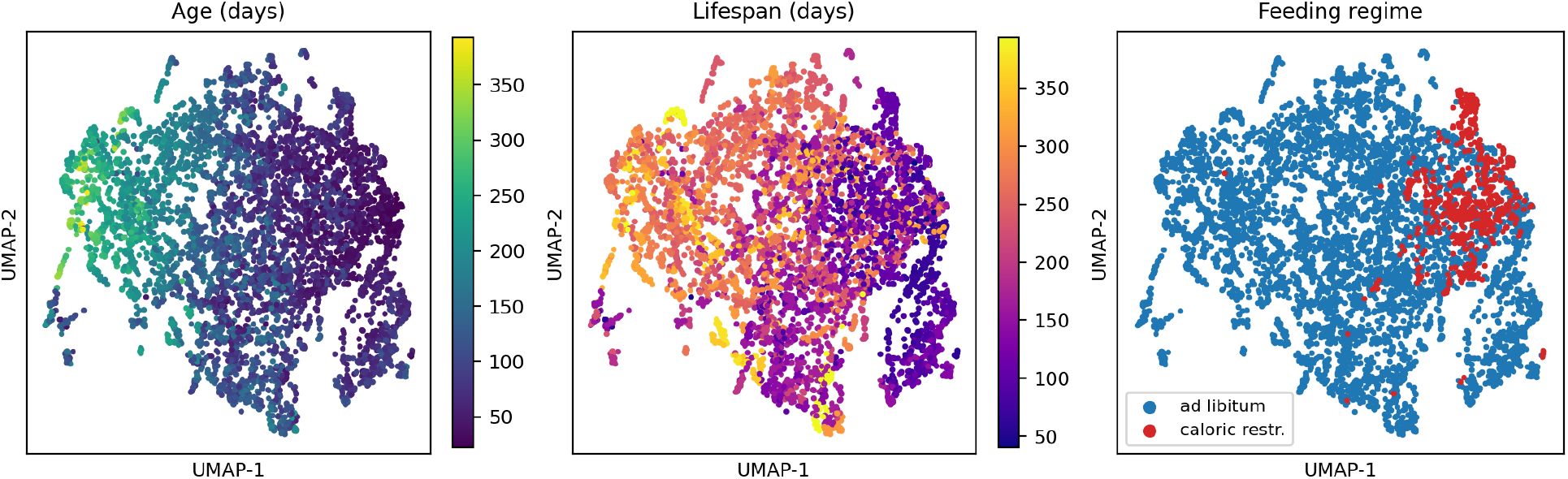
UMAP of day-encoder embeddings (6,000 fish-days, subsampled), colored by age, lifespan, and feeding regime. The representation is organized along biologically meaningful axes: a smooth age gradient, lifespan structure, and partial separation of *ad libitum* from caloric-restriction fish, none of which were used during self-supervised pre-training.

### 4.2 A single day suffices: the life-course stage adds nothing

Holding the day encoder fixed, we train the life-course encoder (with the window-relative age encoding of Section 3.3) and ask whether it adds anything over the day-level representation it is built on. Unless noted, training uses full per-fish coverage. Across all three targets with full-life ground truth the answer is no: each is matched by the single-day representation plus a *trivial, non-learned* operation, with no learned trajectory model needed (Table 3).

**Table 3.**
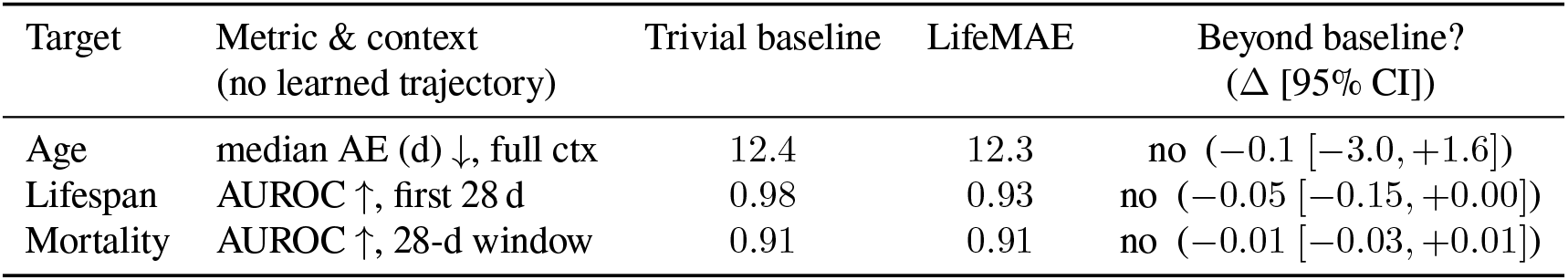
Does the life-course stage beat a trivial, non-learned baseline? For each target: the strongest baseline that uses *no* learned trajectory model: a moving-average smoother of the per-day predictions for age, an early-life mean-pool for lifespan, and the single day plus its within-window position (Section 4.3) for mortality; then LifeMAE (window-relative encoding, full-coverage training), and whether LifeMAE improves on it (Δ = LifeMAE − baseline, signed per the metric’s arrow; 95% fish-level paired bootstrap over a 3 × 3 seed factorial). For none of the three does the life-course stage add measurable value.

| Target | Metric & context<br>(no learned trajectory) | Trivial baseline | LifeMAE | Beyond baseline?<br>($\Delta$ [95% CI]) |
| --- | --- | --- | --- | --- |
| Age | median AE (d) ↓, full ctx | 12.4 | 12.3 | no (−0.1 [−3.0, +1.6]) |
| Lifespan | AUROC ↑, first 28 d | 0.98 | 0.93 | no (−0.05 [−0.15, +0.00]) |
| Mortality | AUROC ↑, 28-d window | 0.91 | 0.91 | no (−0.01 [−0.03, +0.01]) |

#### Age and lifespan are already legible day by day

For chronological age and eventual lifespan the day-level representation already suffices, and the life-course stage contributes only aggregation. A single day predicts age at *R*^2^ = 0.88 (Section 4.1); the life-course encoder’s apparent age refinement at full observation context (14.4 → 12.3 d)^1^ is matched by a simple temporal smoother of the per-day predictions (best ±14-day moving average, 12.4 d),^2^ the two statistically indistinguishable (Δ =− 0.1 d, 95% fish-level CI [− 3.0, +1.6]). Eventual lifespan is the same story: mean-pooling the first 14–28 days of behavior predicts long-versus-short lifespan at AUROC 0.96–0.98, and the life-course fish embedding adds nothing over that pool (ΔAUROC = −0.05 [− 0.15, +0.00]); a learned fish-level `[CLS]` read-out does no better than the mean-pool either (ΔAUROC = −0.05 [−0.13, −0.01], if anything worse). Both are read off day-level content: one day for age, the averaged early-life phenotype for lifespan, leaving the trajectory model only denoising to do. The practical upshot is that these targets need almost no longitudinal data: because the day encoder is trained on single days alone, age is readable from one day and eventual lifespan from a short, simply-averaged early-life window, with no trajectory model and no need to follow individuals across their lives.

#### 30-day mortality looks like the exception, but is not

A single day carries little 30-day-mortality signal on its own (AUROC 0.81), and the life-course encoder appears to supply a large gain: given roughly a month of behavior it raises 30-day-mortality AUROC to 0.90–0.93, +0.09 to +0.11 over the best smoother, with fish-level intervals excluding zero. This is the one place the trajectory model looks essential. It is not. The gain is a *within-window position* cue, supplied by the encoder’s representation of time rather than by behavior: once the day-level probe is given that same position information, the life-course encoder adds nothing (ΔAUROC =− 0.01 [− 0.03, +0.01]; Table 3, mortality row), exactly as for age and lifespan. Section 4.3 establishes this in detail. Mortality thus joins age and lifespan: legible from a single day (here, plus where that day sits in the observation window), with nothing left for a learned trajectory model to add.

### 4.3 The apparent mortality gain is a within-window position artifact

The mortality result of Section 4.2 turns on a single control, which we now justify. The life-course encoder is given each day’s position within the observed window through its window-relative encoding (Section 3.3), and for this cohort that position is itself predictive of mortality. The 30-day label is, by construction, a threshold on remaining lifespan, and for a short-lived fish a centered 28-day window covers most of its life, so its near-death days fall in the *late* part of the window; short-lived fish also contribute most positive cases (79 of the 111 positive rows in this 28-day mortality evaluation, which scores 588 rows across the 21 natural-death test fish, come from fish living under 120 days). Within-window position therefore tracks remaining life. It is a strong predictor on its own: *with no behavioral input at all*, the normalized position of a day in its window predicts 30-day mortality at AUROC 0.71 overall, and 0.94 among short-lived fish.

Table 4 makes the consequence explicit. A single-day probe given behavior alone reaches AUROC 0.81; given behavior plus the within-window position vector the encoder itself receives, it reaches 0.91, matching LifeMAE. The entire life-course gain over a position-blind single day (+0.09) is thus the position cue, not learned trajectory structure: LifeMAE does not improve on behavior + position (ΔAUROC = −0.01 [− 0.03, +0.01]), and adding a window-mean of the day embeddings only widens the gap against LifeMAE.

The same control holds at every observation window (Fig. 3). Read naively, the life-course gain over a position-blind single day rises with the length of the observation window to a plateau, the profile that suggests the trajectory model “needs about a month of context” before it helps. Position alone, with no behavior, follows that curve cell for cell; and once the single-day probe is given the position, the residual is flat at zero across the entire ladder and slightly negative at the longest windows. The apparent context requirement is not trajectory information accruing as the window grows; it is position becoming more separable as the window lengthens.

**Table 4.** The 30-day-mortality gain is a within-window position cue, not behavior. 30-day-mortality AUROC (window-relative encoding, retrospective 28-day context, natural-death test fish, 3 × 3 seeds). Position alone (where each day sits in its window, with no behavioral input) already predicts mortality, and a single-day probe given that position matches LifeMAE; so the life-course gain over a position-blind single day (+0.09) is supplied by position. (Δ = LifeMAE − row; 95% fish-level paired bootstrap.)

| Mortality predictor (identical probe) | AUROC | $\Delta$ to LifeMAE [95% CI] |
| --- | --- | --- |
| Single day, behavior only (raw day embedding) | 0.81 | +0.09 [+0.04, +0.18] |
| Within-window position only, no behavior | 0.71 | — |
| Single day + within-window position | 0.91 | −0.01 [−0.03, +0.01] |
| **LifeMAE** (full life-course embedding) | **0.91** | — |

**Figure 3.**
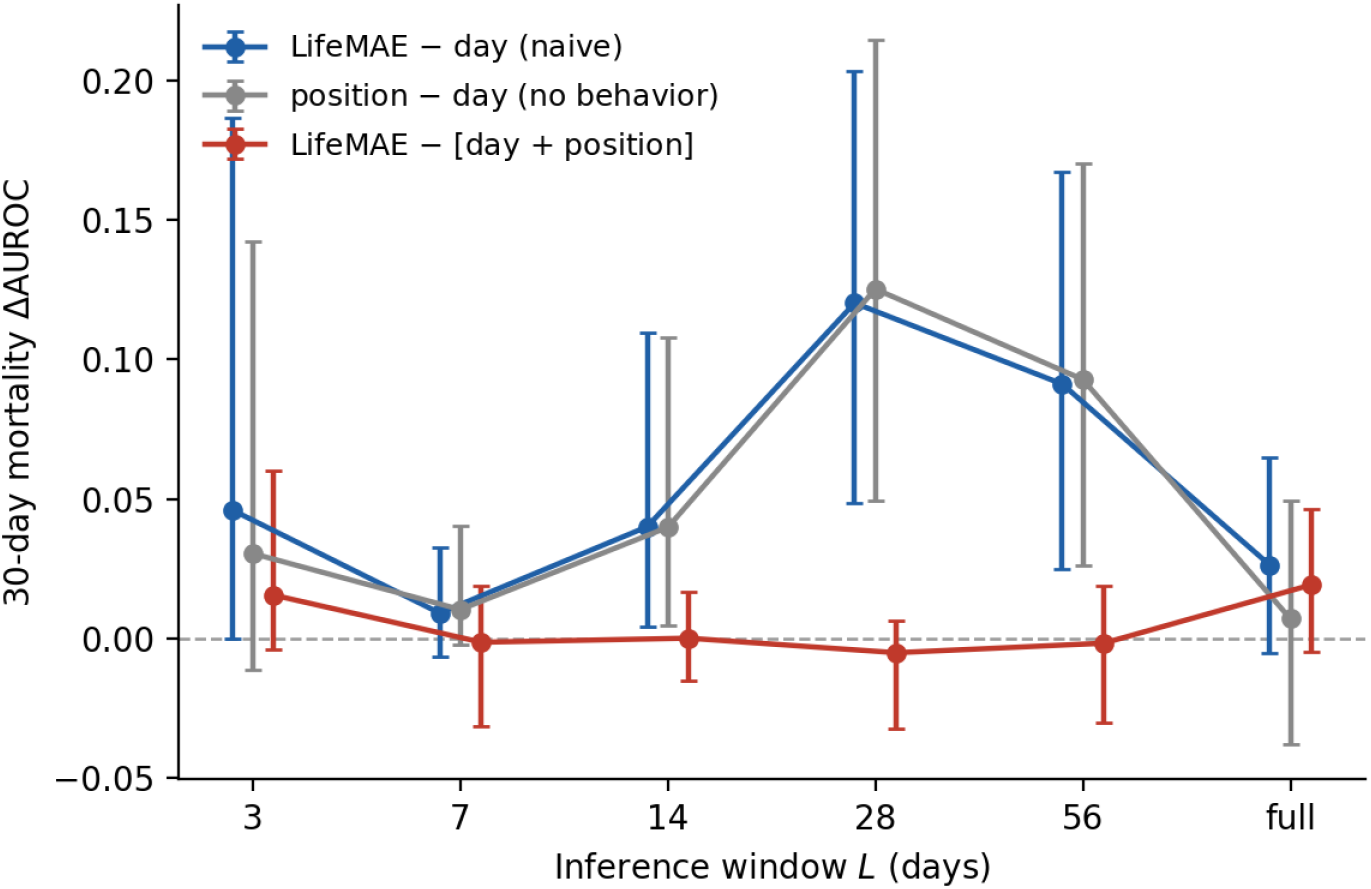
The apparent mortality gain is a within-window position artifact, at every observation window. For each inference-window length *L* (centered, retrospective; full-coverage encoder), the gain of the life-course encoder over a position-blind single day (LifeMAE − day, blue) rises with *L* to a plateau, the profile that, read naively, suggests the trajectory model “needs about a month of context.” Within-window position alone, no behavior (position − day, grey), tracks it cell for cell. Once the single-day probe is given that position (LifeMAE − [day + position], red), the residual is flat at zero across the ladder and slightly negative at the longest windows. Points are the 3 × 3-seed mean; error bars are the 95% fish-level paired bootstrap (*B*=2000). The retired “≥7-day inference floor” is just the blue curve’s rise.

#### This also dissolves the apparent fragmentation tolerance

An encoder trained only on disjoint 3-day episodes reproduces the same apparent mortality gain as one trained on complete lifespans, flat across the training ladder. With the gain identified as a position cue, this robustness is expected and not a capability: the window-relative encoding supplies within-window position regardless of how the encoder was trained, so an apparent “fragmentation-tolerant trajectory signal” is the position artifact, recovered at any training coverage. The fully position-confounded (training × inference) matrix is given in Appendix A.

#### And it is consistent with the causal read-out

Under a *causal* trailing read-out (where the target day is always the last day of the window, so within-window position is fixed and cannot vary with the outcome) the life-course gain is only ∼0.02–0.03 AUROC with fish-level intervals including zero (Section 6). Removing the position degree of freedom removes the effect, as the artifact account predicts.

### 4.4 Evaluation choices that shaped the measurements

The conclusions above rest on a few methodological choices; we record them because they transfer to other longitudinal-representation studies, and because two of them are what separated a real null from an apparent capability.

#### Positional encodings leak task-specific position

A representation will read its positional encoding whenever that encoding correlates with the prediction target, manufacturing apparent skill. We hit this twice, in opposite directions. Encoding each day’s *absolute* age in the life-course positional encoding roughly doubles the apparent age advantage (full-trained: −4.2 vs −2.2 d at full context) and renders it nearly flat across observation context (∼ 4 days whether 7 days or a whole life are seen), the signature of a model recovering chronological age, the prediction target, from its own positional encoding rather than from behavior. Switching to a window-relative encoding removes that age leak. But the window-relative encoding introduces its own: it supplies each day’s *within-window position*, which tracks remaining life and inflates retrospective mortality (Section 4.3). Neither leak is behavioral; both are read off the encoding of time. The general lesson is to treat any positional signal correlated with the target as a confound and control for it directly.

#### Retrospective vs causal read-out

Apparent “minimum-context” thresholds are protocol-specific, and only a read-out fixed to the moment of prediction is free of the position confound. Under a centered (retrospective) read-out a day’s position within the window varies and can track the outcome; under a causal trailing read-out the target day is always at the window edge, so position is fixed. The mortality gain that looks large retrospectively shrinks to ∼0.02–0.03 AUROC, intervals including zero, under the causal read-out (Sections 4.3 and 6). We report both and rest no positive claim on the retrospective protocol alone.

#### Probe expressiveness and multi-seed evaluation

Two further choices mattered just as much. A small nonlinear probe improves the day encoder’s age estimate (median AE 15.7 → 12.6 d) but barely changes the train-only tensor-factorization baseline (20.3 → 19.6 d; Fig. 4), not a generic probe effect but age-relevant structure a linear read-out misses, so a linear-probe-only study would mis-rank the representations. And encoder-initialization variance is comparable to the effects we measure: the day-encoder seed alone accounts for roughly three-quarters of the variance in the life-course gain, single-seed intervals are about half their true width, and single-seed sweeps showed spurious sign flips that dissolved under the 3 × 3 paired bootstrap. We therefore report every comparison multi-seed, with fish-level intervals; at this sample size single-seed evaluation systematically overstates confidence.

**Figure 4.**
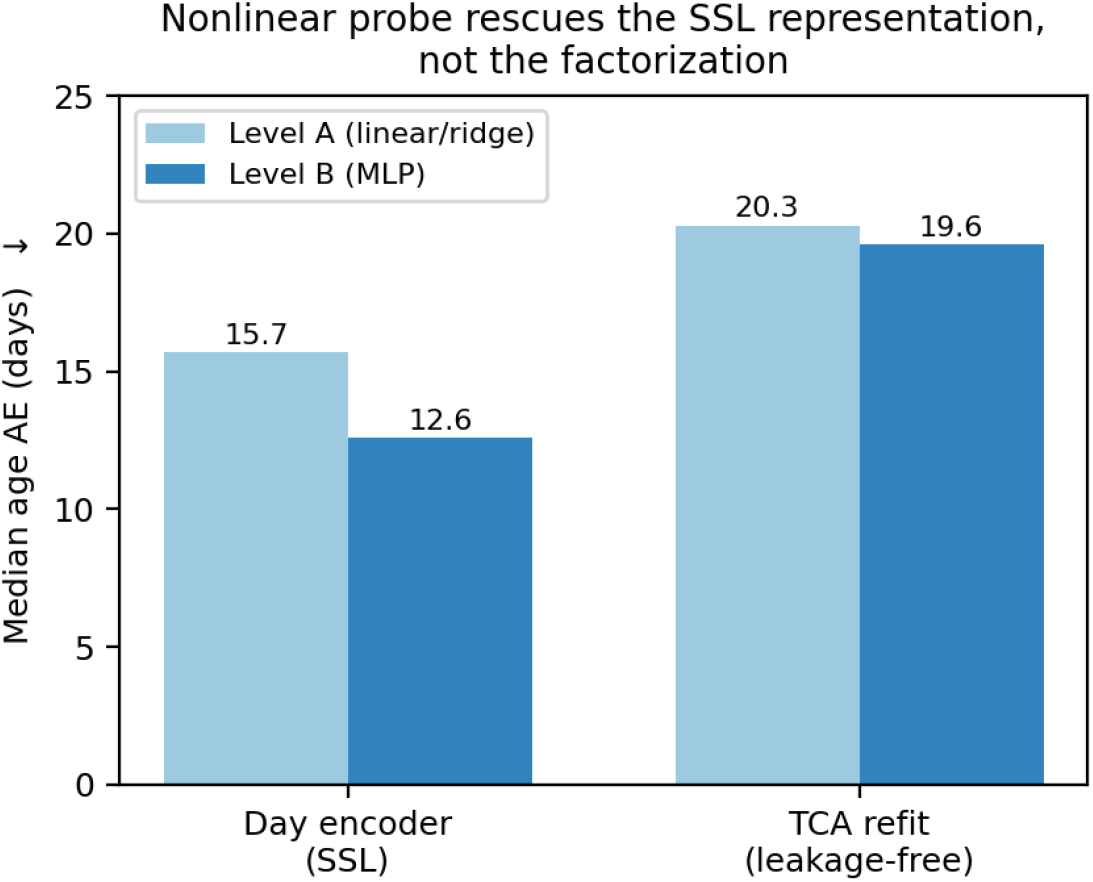
Probe expressiveness is representation-specific. Per-fish-day *median* age error under a linear (Level A) and a nonlinear (Level B) probe: the nonlinear probe improves the self-supervised day encoder by ∼3 days but barely changes the train-only tensor-factorization baseline.

## 5 Discussion

### What modeling the whole life course bought, and did not

For self-supervised models trained on biobank-, electronic-health-record-, or wearable-style data, where each individual contributes only a fragment of a long life, our results give an unusually clean piece of guidance, because we could check against complete lifespans. Across all three traits with full-life ground truth, adding a life-course stage over a single-day representation bought nothing: chronological age and eventual lifespan are already legible from one day (or its average), and the one trait where the trajectory model appeared essential, 30-day mortality, turned out to be reading where each day sat in the observation window, a cue supplied by the model’s encoding of time rather than by behavior (Section 4.3). For these readouts the modeling effort is better spent on the single-day representation than on a trajectory model stacked on top of it. Our negative result joins a growing body of critical benchmarks finding that, under fair comparison, deep and foundation models often fail to outperform deliberately simple baselines; in single-cell transcriptomics, none beat an additive or linear baseline for gene-perturbation prediction [Ahlmann-Eltze et al., 2025]. We reach the same conclusion in a different domain, behavioral aging, and add a specific mechanism for the over-optimism: a positional-encoding leak that lets the complex model read its target off the geometry of the observation window rather than from behavior.

### Why a controlled organism, and what carries

That negative result is quantitative only because the killifish supplies what no human cohort can: complete lifespans for hundreds of individuals, so the life-course stage could be measured against a full-life upper bound rather than against another fragment, and so the apparent mortality gain could be decomposed against ground-truth lifespan and exposed as positional. The operational feasibility of fragmentary longitudinal self-supervision was already established at human scale; what a short-lived vertebrate uniquely provides is the controlled counterfactual that separates a genuine trajectory capability from a positional artifact. Neither the sufficiency of single-day information nor the positional-encoding trap is killifish-specific, and both are directly testable on human longitudinal data.

### Implications for cohort design

These results bear on how to collect longitudinal aging data. For every trait we could ground-truth, the read-out needs at most a single day per individual: age from one day, eventual lifespan from a short early-life window, and near-term mortality from one day plus where it sits in the record. They point the same way, against the instinct to track each individual as long as possible: under a fixed observation budget, breadth (many individuals, each observed briefly) is likely worth more than depth (a few tracked exhaustively) for characterizing an individual’s state. Two caveats keep this a prediction rather than a result. We hold the day encoder fixed; although it is itself trained cross-sectionally, on single days pooled across the cohort, we do not run a controlled breadth-versus-depth experiment at matched budget. And our claim is about *characterizing* present state, not forecasting: a genuinely forward-looking, deployable predictor was weak here (Section 6), and a trait that turns on the *rate* of behavioral change (which we did not find among the three we could test) might still reward a trajectory model. A controlled breadth-versus-depth experiment, and a search for trajectory-dependent targets, are natural next steps.

### Measurement lessons

Two evaluation choices materially changed our conclusions and transfer to other longitudinal-representation studies (Section 4.4). First, a learned representation will read its positional encoding whenever that encoding correlates with the target: an absolute-age encoding let the model recover age, its own target, and a window-relative encoding let it recover within-window position, which tracks remaining life: two leaks in opposite directions, both mistakable for learned biology unless controlled directly, and only the second is removed by fixing the read-out to the moment of prediction. Second, at modest cohort sizes initialization variance rivals the effects of interest, so multi-seed, individual-level intervals are needed to avoid spurious conclusions.

### Scope and what cannot be said

All fish come from a single laboratory cohort of one substrain, processed through one upstream behavioral tokenizer; cross-laboratory and cross-strain generalization is untested, and the representation inherits whatever the upstream hidden Markov model discards (Section 6). We make no cross-species claims, and we do not claim that life-course modeling is useless in general, only that, for three well-grounded aging readouts in this controlled system, it added nothing a single day did not already carry. What we provide is a controlled map of when life-course modeling helps, and a cautionary account of how it can appear to help when it does not: a prerequisite, not a substitute, for the human longitudinal studies that motivate the question.

## 6 Limitations

### The claim is about the trajectory-aggregation stage

Our fragmentation ablation holds the day encoder fixed (trained once on all training fish-days) and fragments only the life-course stage, so our conclusions concern the trajectory-aggregation stage given a fixed day representation. We argue (Section 5) that this matches how fragmentary cohorts are actually structured, but we do not test whether the day encoder itself can be learned under matched fragmentation; an end-to-end fragmented pipeline is left to future work.

### The position-controlled (causal) effect is small and unresolved

The mortality gain that appears under a retrospective read-out is a within-window position cue (Section 4.3); under a causal trailing read-out (where the target day is the last in the window, so position is fixed and cannot track the outcome) it falls to ∼ 0.02– 0.03 AUROC with fish-level intervals including zero, limited by the 116-fish natural-death cohort. Whether this residual is a genuinely small behavioral effect or zero cannot be resolved at this sample size; a larger death cohort (or a human cohort) is needed, and we make no positive deployable-mortality claim.

### We cannot isolate a behavioral decline signal beyond position

Because the 30-day label is a threshold on remaining lifespan and the window-relative encoding makes within-window position available, the apparent life-course mortality gain is accounted for by position alone (Table 4, Fig. 3): a single-day probe given the same position matches LifeMAE. We therefore cannot claim a behavioral terminal-decline signal distinct from where an animal sits in its record; whether such a signal exists (separable from position and from day-level age and lifespan) would need a larger natural-death cohort and a position-free read-out.

### Three traits, one cohort, one pipeline

Our conclusions rest on three readouts with full-life ground truth (chronological age, eventual lifespan, 30-day mortality); whether some other, genuinely trajectory-dependent trait would reward the life-course stage is untested; we simply found none among the three. All 234 fish come from a single laboratory cohort of one *N. furzeri* substrain, recorded and tokenized by one pipeline [Bedbrook et al., 2026]; cross-laboratory and cross-strain generalization is untested. The day encoder also inherits whatever the upstream 100-state hidden-Markov tokenizer discards (continuous within-syllable kinematics, microstructure finer than the 10-minute bin) and the public release includes 20 Hz pose for only 82 of 24,277 fish-days, so a learned pose-level front end is not yet possible.

### Small by deep-learning standards

164 training fish is substantial for longitudinal aging biology but small for representation learning, which is precisely why initialization variance rivals the effects we measure (Section 4.4) and why we evaluate with frozen-embedding probes rather than fine-tuning.

#### Future directions

Several questions follow directly. We do not compare temporal architectures: whether a recurrent or convolutional aggregator, or a generic self-supervised time-series model, matches the Transformer is open. Early-life behavior predicts eventual lifespan well, but at the level of the averaged early phenotype; whether a trajectory model can forecast longevity beyond that, and whether any trait that turns on the *rate* of behavioral change rewards trajectory modeling, are natural extensions, currently limited by per-individual sample size. The natural next tier is to carry the same controlled question to human biobank, wearable, or clinical-record cohorts, using the killifish result as a calibrated reference for which capabilities need the trajectory and how much per-individual observation they require, and as a reminder to control for positional leakage when a trajectory model appears to help.

## 7 Conclusion

Self-supervised models of human aging must learn from fragmentary, partial-life data, yet the field has had no way to ask, in a controlled fashion, which capabilities actually need the longitudinal trajectory, because no human cohort offers full-life ground truth. Using the African turquoise killifish as a controlled testbed, with complete behavioral lifespans for hundreds of individuals, we asked whether a learned life-course representation adds anything over a single day of behavior, and found that it does not: across chronological age, eventual lifespan, and 30-day mortality, a single-day representation plus a trivial, non-learned operation matches the two-stage model. The one trait where the life-course stage appeared essential, near-term mortality, was reading where each day fell within the observation window (a cue introduced by how the model encodes time, not a behavioral trajectory) and the apparent gain vanished once a single-day model was given the same information. For these aging readouts an individual’s place in its life course is legible from a snap-shot, which favors observing many individuals briefly over tracking a few for long. The result also leaves a transportable warning for the longitudinal foundation models this work is meant to inform: the way such models encode time can leak the very quantity being predicted, and ordinary backward-looking evaluation will mistake it for learned biology unless the read-out is fixed to the moment of prediction. Whether some trait that genuinely turns on the rate of behavioral change rewards a trajectory model, and whether these lessons hold on human data, are the questions we would carry forward.

## Data and code availability

The behavioral data analyzed in this study were generated by Bedbrook et al. [2026] and are publicly available from Zenodo (10.5281/zenodo.17238217, CC-BY-4.0); none of the data are redistributed here. Code to reproduce all analyses and figures, together with the derived result files, is available at https://github.com/jenchien/lifemae.

## Declaration on the use of AI

The authors used two AI assistants: Claude (Anthropic), which contributed to the design and implementation of the software pipeline used to train the models and run the analyses in this study, and to the drafting and editing of this manuscript; and ChatGPT (OpenAI), for critical feedback on draft revisions. The conception of the project, all scientific claims, and the verification of the results are the responsibility of the authors.

## Acknowledgements

We thank Takuma Kasai and Steffen Backes (Research DX Foundation Team, RIKEN) for valuable discussions and for reviewing our analysis code.

## Competing interests

The authors declare no competing interests.

## A Full training×inference-context matrix (window-relative encoding)

Table 5 gives the per-cell *apparent* life-course value-add under the window-relative encoding (Section 3.3) and the retrospective read-out, for every (training regime × inference-context) cell, that is, before the within-window position control of Section 4.3. Each entry is the Δ between the life-course encoder and the day-level (per-day) baseline, averaged over the 3 × 3 day-encoder×life-course-encoder seed factorial (point estimate); negative favors the life-course encoder for age (lower MAE), positive for mortality (higher AUROC). Fish-level paired-bootstrap intervals for the headline cells (age at full context, 30-day mortality at the 28-day context) are reported in the main text (Table 3). These are the numbers a naive analysis would report. The main text shows that the age column is matched by a temporal smoother of the per-day predictions (Table 3), i.e. denoising rather than trajectory structure, and that the mortality column is a within-window position cue that collapses to ≈ 0 once position is controlled, at every inference context (Table 4, Fig. 3).

**Table 5.** (Training regime × inference-context) *apparent* value-add, retrospective read-out, window-relative encoding, before the within-window position control of Section 4.3. Columns are the inference-context length *L*_inf_ in days (“full” = the complete observed window); Δ is life-course encoder − day-level baseline, 9-seed mean.

| Training regime | Age ΔMAE (days) ↓ | | | Mortality ΔAUROC ↑ | | |
| --- | --- | --- | --- | --- | --- | --- |
|  | 7 | 28 | full | 7 | 28 | full |
| fixed-3 | −1.72 | −1.13 | −3.25 | 0.00 | +0.12 | +0.09 |
| fixed-7 | −1.70 | −1.05 | −3.28 | 0.00 | +0.13 | +0.10 |
| fixed-14 | −1.34 | −1.92 | −2.67 | 0.00 | +0.12 | +0.10 |
| fixed-28 | −0.72 | −0.97 | −2.31 | 0.00 | +0.11 | +0.09 |
| fixed-56 | −0.67 | −1.23 | −2.52 | −0.01 | +0.10 | +0.07 |
| full | +0.34 | −1.38 | −2.17 | −0.01 | +0.10 | +0.02 |

Two patterns are visible in these uncontrolled numbers, and both are the position artifact rather than a trajectory capability. *Down the columns* (training coverage, at fixed inference context) the apparent value-add is roughly flat (the “fragmentation tolerance” of earlier drafts) because the window-relative encoding supplies within-window position regardless of how much of each life was seen in training. *Across the mortality columns* the apparent gain requires roughly a month of observation (≈ 0 at 7 days, peaking near 28); this is position becoming separable as the window lengthens, not terminal decline accruing. The position-controlled and causal read-outs of the same cells are flat at zero (Fig. 3, Section 6).

## Footnotes

1 This 14.4 d day-level baseline is evaluated inside the life-course harness and averaged over the three day-encoder seeds, so it is paired with the LifeMAE read-out; it runs above the 12.6 d single-seed cohort-wide probe of Table 2 because encoder-initialization variance is large at this cohort size (Section 4.4). The paired difference within each evaluation is what we report.

2 The smoothing width is chosen to maximize the baseline on the evaluation fish (an oracle upper bound on simple smoothing) over ± {1, 3, 7, 14} d for age and ± {1, 3, 7} d for mortality, so each life-course–versus–smoother comparison is conservative toward the baseline.

## Notes

### Competing Interest Statement

The authors have declared no competing interest.

## References

Constantin Ahlmann-Eltze, Wolfgang Huber, and Simon Anders. Deep-learning-based gene perturbation effect prediction does not yet outperform simple linear baselines. Nature Methods, 22:1657–1661, 2025. doi: 10.1038/s41592-025-02772-6.

Mehdi Azabou, Michael Mendelson, Nauman Ahad, Maks Sorokin, Shantanu Thakoor, Carolina Urzay, and Eva L. Dyer. Relax, it doesn’t matter how you get there: A new self-supervised approach for multi-timescale behavior analysis. In NeurIPS, 2023.

Claire N. Bedbrook et al. Lifelong behavioral screen reveals an architecture of vertebrate aging. Science, 391(6790): eaea9795, 2026. doi: 10.1126/science.aea9795. Zenodo data release: https://doi.org/10.5281/zenodo.17238217 (CC-BY-4.0).

Xinlei Chen and Kaiming He. Exploring simple siamese representation learning. CVPR, 2021.

Edward Choi, Zhen Xu, Yujia Li, Michael W. Dusenberry, Gerardo Flores, Yuan Xue, and Andrew M. Dai. Learning the graphical structure of electronic health records with graph convolutional transformer. AAAI, 2020.

Sandeep Robert Datta, David J. Anderson, Kristin Branson, Pietro Perona, and Andrew Leifer. Computational neuroethology: a call to action. Neuron, 104(1):11–24, 2019.

Jacob Devlin, Ming-Wei Chang, Kenton Lee, and Kristina Toutanova. BERT: Pre-training of deep bidirectional Transformers for language understanding. In NAACL-HLT, 2019.

Emadeldeen Eldele, Mohamed Ragab, Zhenghua Chen, Min Wu, Chee Keong Kwoh, Xiaoli Li, and Cuntai Guan. Time-series representation learning via temporal and contextual contrasting. In IJCAI, 2021.

Itamar Harel, Bérénice A. Benayoun, Benjamin E. Machado, Param Priya Singh, Chi-Kuo Hu, Matthew F. Pech, Dario R. Valenzano, Elisa Zhang, Sabrina C. Sharp, Steven E. Artandi, and Anne Brunet. A platform for rapid exploration of aging and diseases in a naturally short-lived vertebrate. Cell, 160(5):1013–1026, 2015.

Kaiming He, Xinlei Chen, Saining Xie, Yanghao Li, Piotr Dollár, and Ross Girshick. Masked autoencoders are scalable vision learners. In CVPR, 2022.

Mandar Joshi, Danqi Chen, Yinhan Liu, Daniel S. Weld, Luke Zettlemoyer, and Omer Levy. SpanBERT: Improving pre-training by representing and predicting spans. TACL, 8, 2020.

Yikuan Li et al. BEHRT: Transformer for electronic health records. Scientific Reports, 10:7155, 2020.

Kevin Luxem, Petra Mocellin, Falko Fuhrmann, Johannes Kurzinger, Stephanie R. Miller, Jens F. Sauer, Matthew R. Whiteway, Stefan Remy, and Pavol Bauer. Identifying behavioral structure from deep variational embeddings of animal motion. Communications Biology, 5:1267, 2022.

Alexander Mathis, Pranav Mamidanna, Kevin M. Cury, Taiga Abe, Venkatesh N. Murthy, Mackenzie W. Mathis, and Matthias Bethge. DeepLabCut: markerless pose estimation of user-defined body parts with deep learning. Nature Neuroscience, 21:1281–1289, 2018.

Yuqi Nie, Nam H Nguyen, Phanwadee Sinthong, and Jayant Kalagnanam. A time series is worth 64 words: Long-term forecasting with Transformers. In ICLR, 2023.

Talmo D. Pereira, Nathaniel Tabris, Arie Matsliah, David M. Turner, Junyu Li, Shruthi Ravindranath, Eleni S. Papadoyannis, Edna Normand, David S. Deutsch, Z. Yan Wang, et al. SLEAP: A deep learning system for multi-animal pose tracking. Nature Methods, 19:486–495, 2022.

Laila Rasmy, Yang Xiang, Ziqian Xie, Cui Tao, and Degui Zhi. Med-BERT: pretrained contextualized embeddings on large-scale structured electronic health records for disease prediction. npj Digital Medicine, 4:86, 2021.

Steffen Schneider, Jin Hwa Lee, and Mackenzie Weygandt Mathis. Learnable latent embeddings for joint behavioural and neural analysis. Nature, 617:360–368, 2023. doi: 10.1038/s41586-023-06031-6.

Artem Shmatko et al. Learning the natural history of human disease with generative transformers. Nature, 647:248–256, 2025. doi: 10.1038/s41586-025-09529-3.

Lucas Stoffl et al. Elucidating the hierarchical nature of behavior with masked autoencoders. In European Conference on Computer Vision (ECCV), 2024.

Nicholas Stroustrup, Winston E. Anthony, Zachary M. Nash, Vivek Gowda, Adam Gomez, Isaac F. López-Moyado, Javier Apfeld, and Walter Fontana. The temporal scaling of caenorhabditis elegans ageing. Nature, 530:103–107, 2016.

Kai Wang, Fei Liu, Wei Wu, Changxi Hu, Xian Shen, Meihao Wang, Gen Li, Fanxin Zeng, Li Liu, Io Nam Wong, et al. A full life cycle biological clock based on routine clinical data and its impact in health and diseases. Nature Medicine, 31:4225–4235, 2025a. doi: 10.1038/s41591-025-04006-w.

Yanchen Wang et al. Animal behavioral analysis and neural encoding with transformer-based self-supervised pretraining. arXiv preprint arXiv:2507.09513, 2025b.

Caleb Weinreb, Jonah E. Pearl, Sherry Lin, Mohammed Abdal Monium Osman, Libby Zhang, Sidharth Annapragada, Eli Conlin, Red Hoffmann, Sofia Makowska, Winthrop F. Gillis, et al. Keypoint-MoSeq: parsing behavior by linking point tracking to pose dynamics. Nature Methods, 21:1329–1339, 2024.

Alex H. Williams, Tony Hyun Kim, Forea Wang, Saurabh Vyas, Stephen I. Ryu, Krishna V. Shenoy, Mark Schnitzer, Tamara G. Kolda, and Surya Ganguli. Unsupervised discovery of demixed, low-dimensional neural dynamics across multiple timescales through tensor component analysis. Neuron, 98(6):1099–1115, 2018.

Alexander B. Wiltschko, Matthew J. Johnson, Giuliano Iurilli, Ralph E. Peterson, Jesse M. Katon, Stan L. Pashkovski, Victoria E. Abraira, Ryan P. Adams, and Sandeep Robert Datta. Mapping sub-second structure in mouse behavior. Neuron, 88(6):1121–1135, 2015.

Zhenda Xie, Zheng Zhang, Yue Cao, Yutong Lin, Jianmin Bao, Zhuliang Yao, Qi Dai, and Han Hu. SimMIM: A simple framework for masked image modeling. In CVPR, 2022.

Maxwell A. Xu et al. RelCon: Relative contrastive learning for a motion foundation model for wearable data. In International Conference on Learning Representations (ICLR), 2025.

Hang Yuan, Shing Chan, Andrew P. Creagh, et al. Self-supervised learning for human activity recognition using 700,000 person-days of wearable data. npj Digital Medicine, 7:91, 2024. doi: 10.1038/s41746-024-01062-3.

Zhihan Yue, Yujing Wang, Juanyong Duan, Tianmeng Yang, Congrui Huang, Yunhai Tong, and Bixiong Xu. TS2Vec: Towards universal representation of time series. In AAAI, 2022.

George Zerveas, Srideepika Jayaraman, Dhaval Patel, Anuradha Bhamidipaty, and Carsten Eickhoff. A Transformer-based framework for multivariate time series representation learning. In KDD, 2021.

